# Characterization of a spiramycin-inactivating macrolide esterase

**DOI:** 10.64898/2026.10.26.761979

**Authors:** Iryna Myziuk, Zahi Mulla, Taslimun Jannat, Antonio Ruzzini

## Abstract

Recent functional annotation of α/β-hydrolases as macrolide esterases has the opened door to genome mining efforts exploring the sequencing diversity of the family. Nevertheless, unexplained resistance phenotypes associated with these enzymes remain in the literature. Inspired by a report of macrolide-resistant *Streptococcus uberis*(1), we have annotated and characterized an enzyme named EstS as a macrolide esterase. Curiously, phylogenetic analyses revealed that EstS is most often encoded by *Escherichia coli*, including human and animal pathogens, rather than Gram-positive organisms. Although EstS could hydrolyze a series of macrolides, heterologous expression of *estS* conferred protection against only the natural products spiramycin and tylosin. Steady-state kinetic analyses by isothermal titration calorimetry and high-performance chromatography coupled to mass spectrometry revealed that EstS prefers spiramycin as a substrate consistent with higher protection observed in *E. coli*.

---

Bacterial resistance to macrolides through enzymatic hydrolysis was established in the 1980s with the discovery of the first erythromycin esterase(2, 3). Similarly, an α/β hydrolase named EstT, which belongs to a disparate enzyme family than the erythromycin esterases, was reported to hydrolyze veterinary macrolides containing 16-instead of 14-atom macrocycles in 2023(4). EstT-like enzymes are widespread in nature, constituting a group with remarkable sequence diversity that act on 16-atom-containing macrocyclic lactones. Biochemical and biophysical studies of EstT and homologs revealed promiscuous macrolide binding but much more specific hydrolysis with tylosin as a preferred substrate(5). Genome-mining efforts rather than phenotypic screening of bacteria for EstT-like enzymes are ongoing and driving additional in vitro annotations(6-10).

Phenotypic work that predates identification of EstT, namely macrolide resistance in *Streptococcus uberis* causing mastitis in cattle(1), prompted our investigation of a *rdmC*-like gene as a spiramycin resistance determinant. Many α/β-hydrolase gene (*abh*) sequences have been transfer annotated as *rdmC*-like; however, they co-occur with known antimicrobial resistance (AMR) genes (ARGs) and have low sequence identities to *rdmC*, which encodes an aclacinomycin 10-methylesterase(11). An *S. uberis* UCN60 *rdmC*-like gene co-occurs with a macrolide 2’-phosphotransferase II (MPH(2’)II) named *mphB* (Figure 1A). Heterologous expression of both the *mphB* and *rdmC*-like gene were shown to inactivate spiramycin, though the mechanism of the latter was not studied(1). We hypothesised that this *rdmC*-like gene encodes a macrolide esterase with specificity for spiramycin.

**Figure 1.**
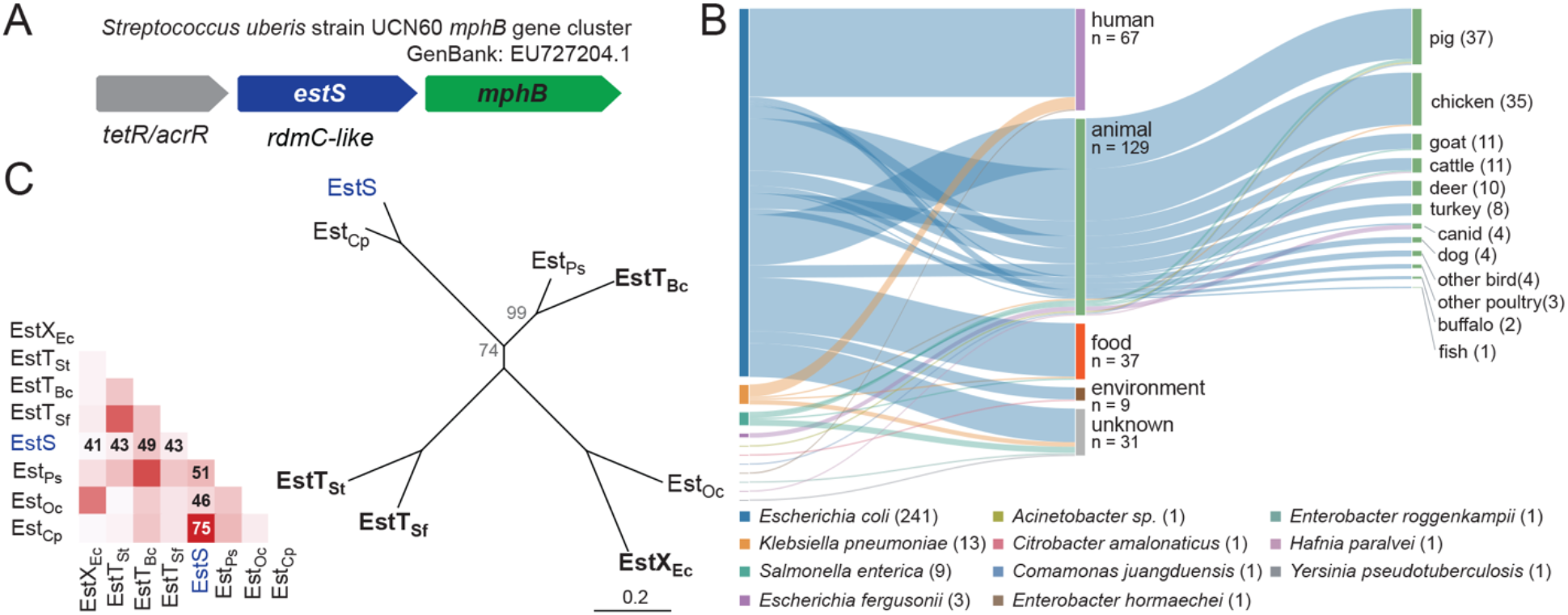
Bioinformatic analysis of an *rdmC*-like gene. **(A)** Gene map of the *S. uberis* UCN60 *mphB* gene cluster that encodes an *rdmC*-like gene renamed *estS*. **(B)** Sankey plot showing the distribution of the *rdmC*-like gene from EU727204.1 in bacteria and corresponding origin of bacterial isolates. **(C)** Phylogenetic tree and heat map showing relatedness of macrolide esterases. Bold font indicates those that have been structurally characterized(5). Accession numbers for the sequences used for the alignment are as follows EstS (ACE78319.1), EstT_Sf_ (UXD71803.1), EstX_Ec_ (PP210101.1), EstT_Bc_ (MFC8766929.1), EstT_St_ (WP_028071475.1), Est_Cp_ (YAI77267.1), EstPs (PQ623322.1), EstOc (MBF2028279.1).

A query of the *S. uberis* UCN60 *rdmC*-like gene product (Genbank no. ACE78319.1; NCBI RefSeq database) revealed matching sequences that were commonly host-associated, including humans (24%) and animals (47%; Figure 1B). It was most frequently found in *Escherichia coli* accounting for 88% of the 273 matched entries. Indeed, the same *rdmC*-like gene was known to be encoded by *E. coli* plasmids prior to the observation in *S. uberis*(12-14). A series of pathogens were amongst the matched sequences, including *E. coli* and *Klebsiella pneumoniae* causing human bloodstream infections(15, 16) as well as infections of broilers(17) and swine(18). Often, the gene was sequenced due to interests in plasmids bearing known ARGs such as the colistin resistance determinant *mcr-1*(19) or β-lactamases(20, 21) rather than those implicated in spiramycin resistance. All but the *S. uberis* UCN60 query were encoded by Gram-negative bacteria, though the next closest homologs were found in Gram-positive genera such as *Clostridia* (YAI77267.1), *Listeria* (XOG14907.1) and *Staphylococcus* (YAI77271.1). Finally, phylogenetic analysis revealed that EstS is between 40 to 50% identical to functionally annotated macrolide esterases (Figure 1C).

To investigate the function of EstS, we produced and purified a C-terminally His_6_-tagged enzyme as described for EstT(4). Incubation of the enzyme (1 μM) with a series of macrolides (200 μM) in 100 mM potassium phosphate (KPi), pH 7.4, revealed hydrolysis of tylosin (TYL), tylvalosin (TYV), spiramycin (SPM), josamycin (JOS) and kitasamycin (KIT) but not tilmicosin (TIL) or erythromycin (ERY). Losses of antibiotic activity were observed by spotting enzyme-treated macrolides on a lawn *Micrococcus luteus* ATCC 4698 (Figure 2A), and qualitative assessment of hydrolysis was performed by time-dependent monitoring of reactions by high-performance chromatography coupled to mass spectrometry (HPLC/MS; Figure 2B,C). In contrast to these biochemical experiments, constitutive expression of *estS* using an insulated promoter(5, 22) in the hyperpermeable and efflux-deficient *E. coli* BW25113 Δ*bamB*Δ*tolC*(23, 24) revealed protection against only TYL and SPM (Figure 1D), consistent with previous work(1). Minimum inhibitory concentrations (MICs) of 16 μg/mL for both macrolides were measured, which were 4 to 8-fold greater than those measured for control transformants carrying an erythromycin esterase (*ereA*) or an *E. coli* codon-optimised version of the *Streptomyces purpurascens rdmC*. As quality control for the experiments, transformants carrying *estT* demonstrated expected resistance to TYL, TVY(5, 6). Thus, *estS* encodes for a macrolide esterase that can hydrolyze TYL and SPM on a timescale that promotes survival of *E. coli*.

**Figure 2.**
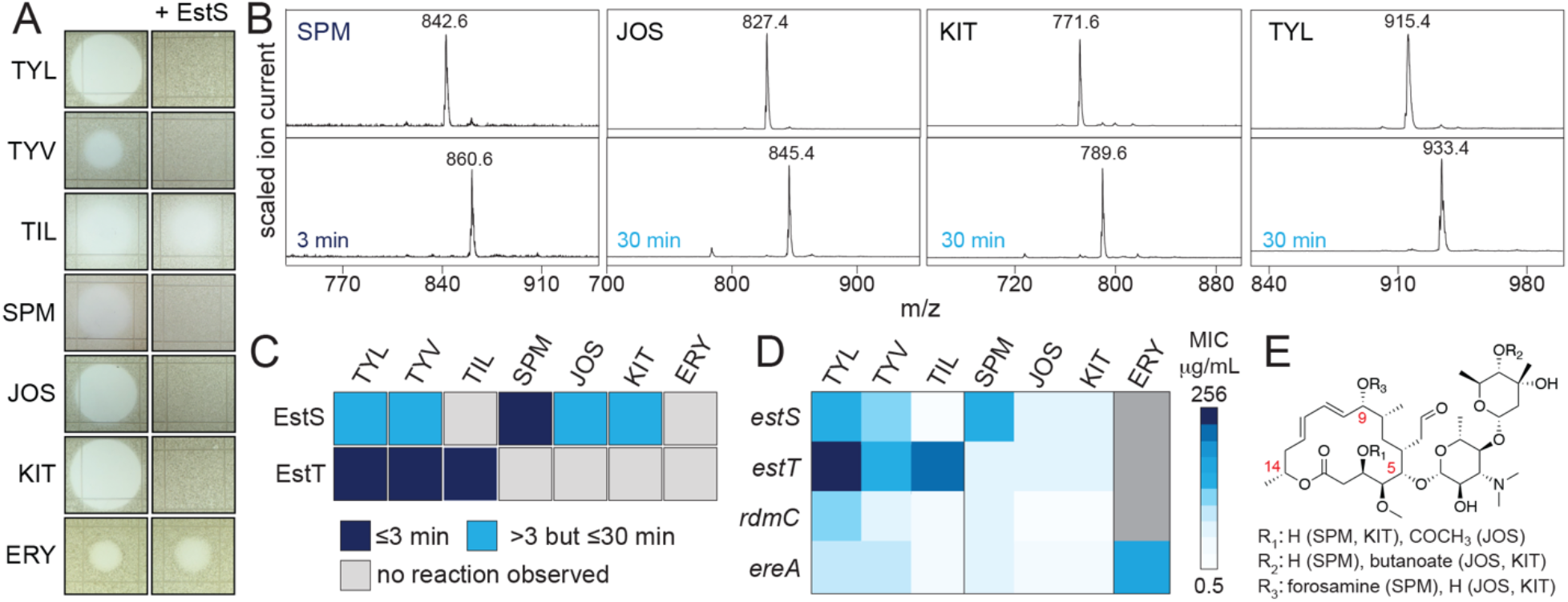
Qualitative analyses of EstL as a macrolide esterase. (A) Antibiotic inactivation shown by spot-on-lawn assays using *M. luteus* as a test strain. (B) Representative mass spectra of EstS-catalysed hydrolysis of SPM, JOS, KIT and TYL. (C) Summary of EstS activity and comparison to EstT, showing macrolides that are hydrolyzed or not. (D) MICs of macrolides determined using *E. coli* 25113 Δ*bamB*Δ*tolC*. (E) Representative macrolide structure showing tailoring specific to SPM, JOS and KIT.

We next sought to establish the substrate specificity of EstS. TYL and SPM were prioritised for study based on their ability to protect *E. coli*. We performed single injection ITC experiments in 100 mM KPi, pH 7.4, to determine steady-state kinetic reaction parameters, following an approach described for EstT and EstX(5). This continuous assay afforded reliable measurements for reactions between EstS and SPM but not TYL (Figure 3A). Although ITC thermograms indicated a reaction between EstS and TYL, the enzyme could not be saturated. In contrast, thermograms of the reaction between 1.5 μM EstS and 500 μM SPM showed a brief exothermic process (dilution of DMSO) that was followed by an endothermic plateau. The latter indicated that the enzyme was saturated with substrate, and the following power inputs represented substrate concentration-dependent changes to the rate of the reaction(25, 26). Steady-state kinetic parameters for the reaction of EstS and SPM were determined to be: *k*_cat_ = 1.07 ± 0.06, *K*_M_ = 127 ± 15, *k*_cat_/*K*_M_ = 8 ± 1 × 10^3^ (Figure 3B). To validate the result, we determined an apparent *k*_cat_ of 1.7 ± 0.3 s^-1^ for the reaction using a discontinuous HPLC/MS assay that relied on quantification of hydrolyzed products(5). The same approach allowed us to determine a specific activity for the reaction of 1 μM EstS and 400 μM TYL to be 0.8 ± 0.3 μM TYL/s^-1^/μM enzyme (Figure 3C). In the broader context of macrolide esterases, the second-order rate constant for the reaction of EstS and SPM is on the same order of magnitude as that observed for EreA and ERY(27) and ∼8-to 42-fold less specific than those determined for EstT_Sf_ and EstX_Ec_ and their cognate substrate TYL.

**Figure 3.**
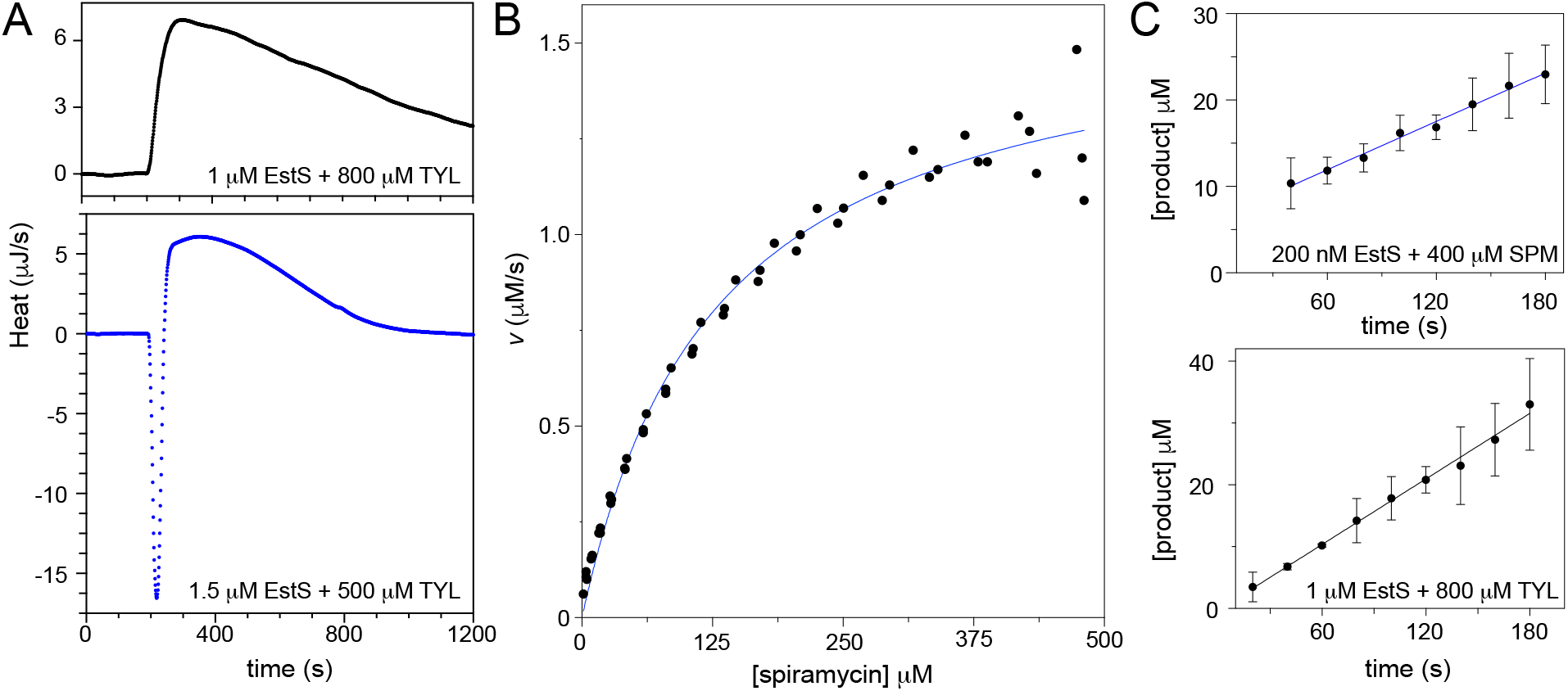
Quantitative characterisation of EstS. **(A)** Representative ITC thermograms of reactions between EstS and TYL (top panel) or SPM (bottom panel). Concentrations of enzymes and substrates are inset. **(B)** Michaelis-Menten plot of the reaction of EstS and SPM generated from three independent ITC measurements. **(C)** HPLC/MS determination of the specific activity of EstS as a tylosin esterase (top panel) and apparent *k*_cat_ for the reaction of EstS and SPM.

Our results call for more robust characterisation of this expanding group of enzymes to ensure accurate functional annotation. Here we establish that EstS can hydrolyze many macrolides in vitro but may only protect against a subset in nature. Macrolide esterases belonging to the α/β-hydrolase superfamily vary greatly in sequence identity (e.g., EstS and EstX: 41%) but less so in substrate specificity with most appearing to prefer TYL. Thus, EstS differs from previously characterized macrolide esterases in that it appears more promiscuous in vitro and is more specific for SPM than TYL. The activity of EstS raises questions about how the α/β-hydrolases have and continue to evolve their roles in antibiotic inactivation. The apparent prevalence of macrolide esterases in *E. coli*, including EstS and EstX, remains puzzling as these organisms are largely considered to be intrinsically resistant to the macrolides they are able hydrolyze.

## ACKNOWLEDGMENTS

Access to the ITC instrument was provided by the Protein Characterization and Crystallization Facility at the University of Saskatchewan and a mass spectrometer provided by PerkinElmer and managed by the Saskatchewan Structural Sciences Centre was used for this study. We thank Gerry Wright (McMaster University) for sharing the *E. coli* BW25113 Δ*bamB*Δ*tolC* strain. This work was supported by a Canadian Institutes of Health Sciences Project Grant to AR and IM is supported by an NSERC CGS-D award.

## Notes

### Competing Interest Statement

The authors have declared no competing interest.

## REFERENCES

1. Achard A, Guerin-Faublee V, Pichereau V, Villers C, Leclercq R. 2008. Emergence of macrolide resistance gene mph(B) in Streptococcus uberis and cooperative effects with rdmC-like gene. Antimicrob Agents Chemother 52:2767–70.

2. Barthelemy P, Autissier D, Gerbaud G, Courvalin P. 1984. Enzymic hydrolysis of erythromycin by a strain of Escherichia coli. A new mechanism of resistance. J Antibiot (Tokyo) 37:1692–6.

3. Ounissi H, Courvalin P. 1985. Nucleotide sequence of the gene ereA encoding the erythromycin esterase in Escherichia coli. Gene 35:271–8.

4. Dhindwal P, Thompson C, Kos D, Planedin K, Jain R, Jelinski M, Ruzzini A. 2023. A neglected and emerging antimicrobial resistance gene encodes for a serine-dependent macrolide esterase. Proc Natl Acad Sci U S A 120:e2219827120.

5. Kelly ETR, Myziuk I, Hemmings MZ, Mulla Z, Blanchet J, Ruzzini A, Berghuis AM. 2026. Hydration and hydrolysis define antibiotic resistance conferred by macrolide esterases. Proc Natl Acad Sci U S A 123:e2611077123.

6. Dhindwal P, Myziuk I, Ruzzini A. 2023. Macrolide esterases: current threats and opportunities. Trends Microbiol 31:1199–1201.

7. Lin J, Lv H, Wang T, Tao H, Zhong Y, Zhou Y, Tang Y, Xie F, Zhuang G, Xu C, Chu Y, Wang X, Yang Y, Song T. 2024. The global distribution of the macrolide esterase EstX from the alpha/beta hydrolase superfamily. Commun Biol 7:781.

8. Zhou Y, Yang Y, Mao Y, Hou Z, Xu Y, Zhao K, Chu Y, Wang X, Wang C, Li S, Xu F, Hao L, Xie B, Lin J, Song T. 2025. Global distribution of alpha/beta hydrolase family macrolide esterases in Gram-positive bacteria. ISME J 19.

9. Tao H, Zhou L, Zhou Y, Wang Y, Lv H, Wang T, Xu C, Chu Y, Wang X, Song T, Lin J. 2026. Functional characterization of macrolide esterase from cyanobacteria and their potential dissemination risk. NPJ Antimicrob Resist 4:10.

10. Song T, Ye X, He Y, Yang T, Zhou J, Chu Y, Wang X, Xu C, Wang W, Lin J. 2026. Functional characterization of the Est-type macrolide esterase ECO-1 from Escherichia coli. Scientific Reports doi:10.1038/s41598-026-62291-y.

11. Niemi J, Mantsala P. 1995. Nucleotide sequences and expression of genes from Streptomyces purpurascens that cause the production of new anthracyclines in Streptomyces galilaeus. J Bacteriol 177:2942–5.

12. Katayama J, Okada H, O’Hara K, Noguchi N. 1998. Isolation and characterization of two plasmids that mediate macrolide resistance in Escherichia coli: transferability and molecular properties. Biol Pharm Bull 21:326–9.

13. Katayama J, Noguchi N. 1999. Nucleotide sequence of the gene cluster containing the mphB gene for macrolide 2’-phosphotransferase II. Biol Pharm Bull 22:227–8.

14. Noguchi N, Katayama J, Sasatsu M. 2000. A transposon carrying the gene mphB for macrolide 2’-phosphotransferase II. FEMS Microbiol Lett 192:175–8.

15. Gladstone RA, McNally A, Pontinen AK, Tonkin-Hill G, Lees JA, Skyten K, Cleon F, Christensen MOK, Haldorsen BC, Bye KK, Gammelsrud KW, Hjetland R, Kummel A, Larsen HE, Lindemann PC, Lohr IH, Marvik A, Nilsen E, Noer MT, Simonsen GS, Steinbakk M, Tofteland S, Vattoy M, Bentley SD, Croucher NJ, Parkhill J, Johnsen PJ, Samuelsen O, Corander J. 2021. Emergence and dissemination of antimicrobial resistance in Escherichia coli causing bloodstream infections in Norway in 2002-17: a nationwide, longitudinal, microbial population genomic study. Lancet Microbe 2:e331–e341.

16. Rigatou A, Afolayan AO, Tatsi EB, Deliolanis I, Michos A, Reuter S, Daikos GL. 2025. Double carbapenemases in Klebsiella pneumoniae blood isolates: dissemination in a single medical center via multiple plasmids and a variety of highly efficient clones. Antimicrob Agents Chemother 69:e0146224.

17. Johnson TJ, Jordan D, Kariyawasam S, Stell AL, Bell NP, Wannemuehler YM, Alarcon CF, Li G, Tivendale KA, Logue CM, Nolan LK. 2010. Sequence analysis and characterization of a transferable hybrid plasmid encoding multidrug resistance and enabling zoonotic potential for extraintestinal Escherichia coli. Infect Immun 78:1931–42.

18. Vereecke N, Van Hoorde S, Sperling D, Theuns S, Devriendt B, Cox E. 2023. Virotyping and genetic antimicrobial susceptibility testing of porcine ETEC/STEC strains and associated plasmid types. Front Microbiol 14:1139312.

19. Martino F, Petroni A, Menocal MA, Corso A, Melano R, Faccone D. 2024. New insights on mcr-1-harboring plasmids from human clinical Escherichia coli isolates. PLoS One 19:e0294820.

20. Garcia-Graells C, Berbers B, Verhaegen B, Vanneste K, Marchal K, Roosens NHC, Botteldoorn N, De Keersmaecker SCJ. 2020. First detection of a plasmid located carbapenem resistant bla(VIM-1) gene in E. coli isolated from meat products at retail in Belgium in 2015. Int J Food Microbiol 324:108624.

21. Matlock W, Rodger G, Pritchard E, Colpus M, Kapel N, Barrett L, Morgan M, Oakley S, Hopkins KL, Roohi A, Karageorgopoulos D, Avison MB, Walker AS, Lipworth S, Stoesser N. 2025. Escherichia coli phylogeny drives co-amoxiclav resistance through variable expression of TEM-1 beta-lactamase. Nat Commun 16:8669.

22. Davis JH, Rubin AJ, Sauer RT. 2011. Design, construction and characterization of a set of insulated bacterial promoters. Nucleic Acids Res 39:1131–41.

23. King AM, Reid-Yu SA, Wang W, King DT, De Pascale G, Strynadka NC, Walsh TR, Coombes BK, Wright GD. 2014. Aspergillomarasmine A overcomes metallo-beta-lactamase antibiotic resistance. Nature 510:503–6.

24. Cox G, Sieron A, King AM, De Pascale G, Pawlowski AC, Koteva K, Wright GD. 2017. A Common Platform for Antibiotic Dereplication and Adjuvant Discovery. Cell Chem Biol 24:98–109.

25. Transtrum MK, Hansen LD, Quinn C. 2015. Enzyme kinetics determined by single-injection isothermal titration calorimetry. Methods 76:194–200.

26. Quinn CF, Hansen LD. 2019. Enzyme Kinetics Determined by Single-Injection Isothermal Titration Calorimetry. Methods Mol Biol 1964:241–249.

27. Morar M, Pengelly K, Koteva K, Wright GD. 2012. Mechanism and diversity of the erythromycin esterase family of enzymes. Biochemistry 51:1740–51.

